## Supplementary Figures for "Molecular characterization of gustatory second-order neurons reveals integrative mechanisms of gustatory and metabolic information"

### Supplementary Figure 1.1

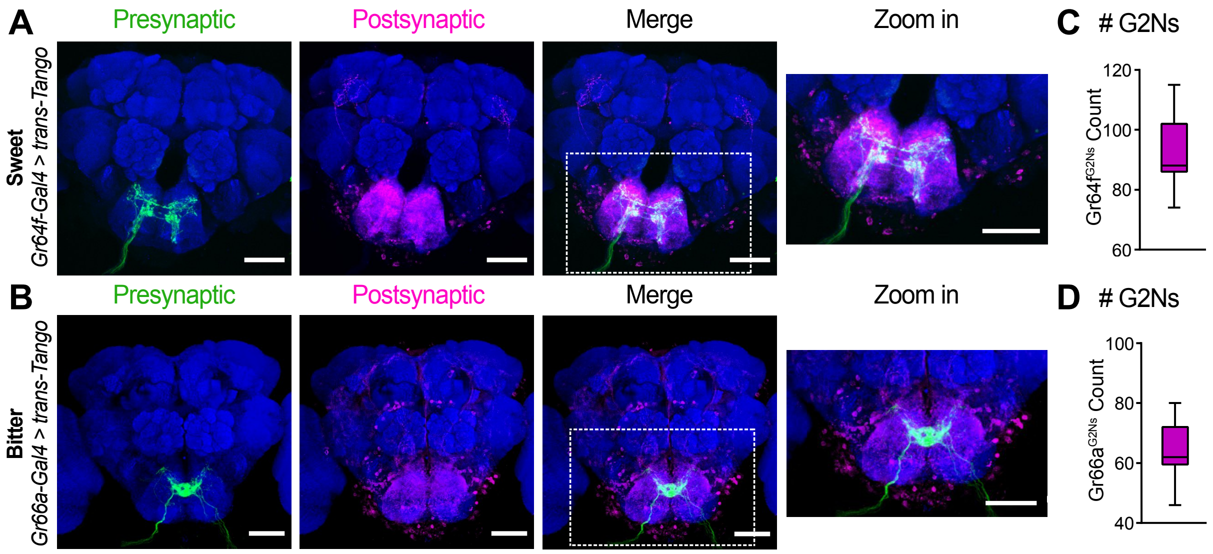


**Supplementary Figure 1.1**. **Variation in G2N numbers for the different populations under study**. Immunofluorescence with anti-GFP (green), anti-RFP (magenta), and anti-nc82 (blue) on a whole-mount brain on a (**A**) *Gr64f-Gal4 > trans-Tango* and (**B**) *Gr66-Gal4 > trans-Tango*. Quantification of the number of G2Ns for each population, (**C**) Gr64f$^{G2Ns}$ (n=11 brains) and (**D**) Gr66a$^{G2Ns}$ (n=9 brains). Larvae were grown at 25°C and 21 days at 18°C after pupation. Scale bar = 50 μm.

**Supplementary Figure 1.2**

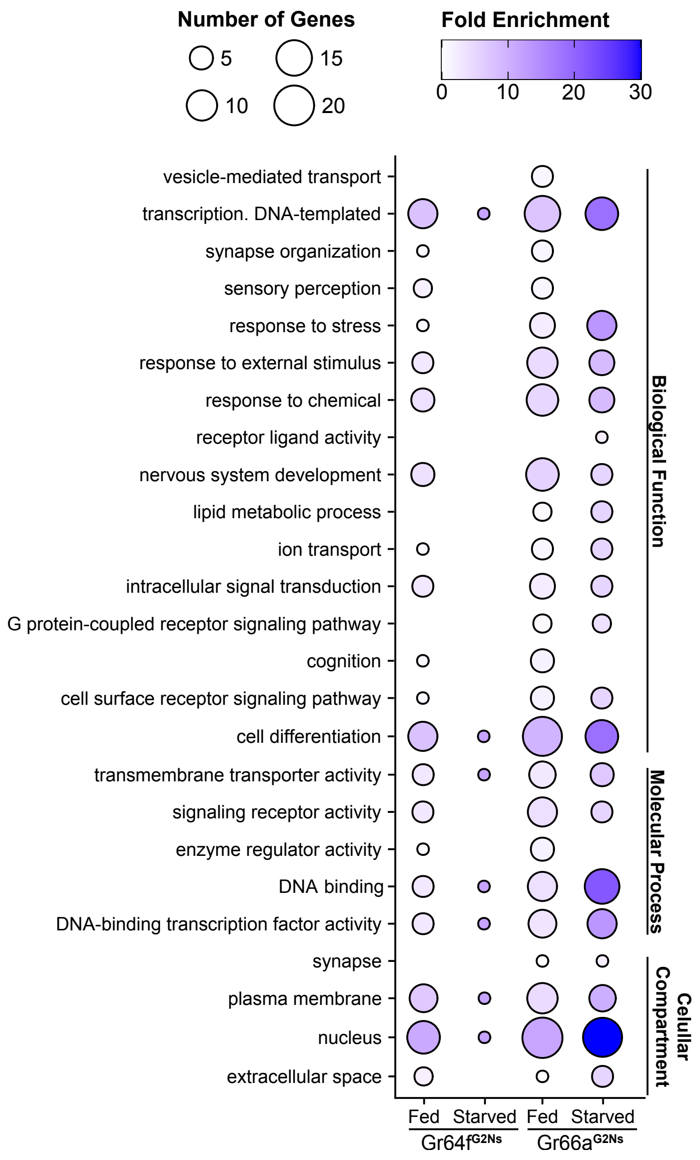


**Supplementary Figure 1.2. Gene Ontology (GO) analysis of the genes differentially expressed in fed and starved conditions for the two G2Ns populations analyzed**. Graphical representation of the GO terms obtained from the GSEA, grouped by the GO category. The data represent the fold enrichment with a heatmap, and the number of genes included in each GO term by size for all G2Ns and metabolic states analyzed.

**Supplementary Figure 6**

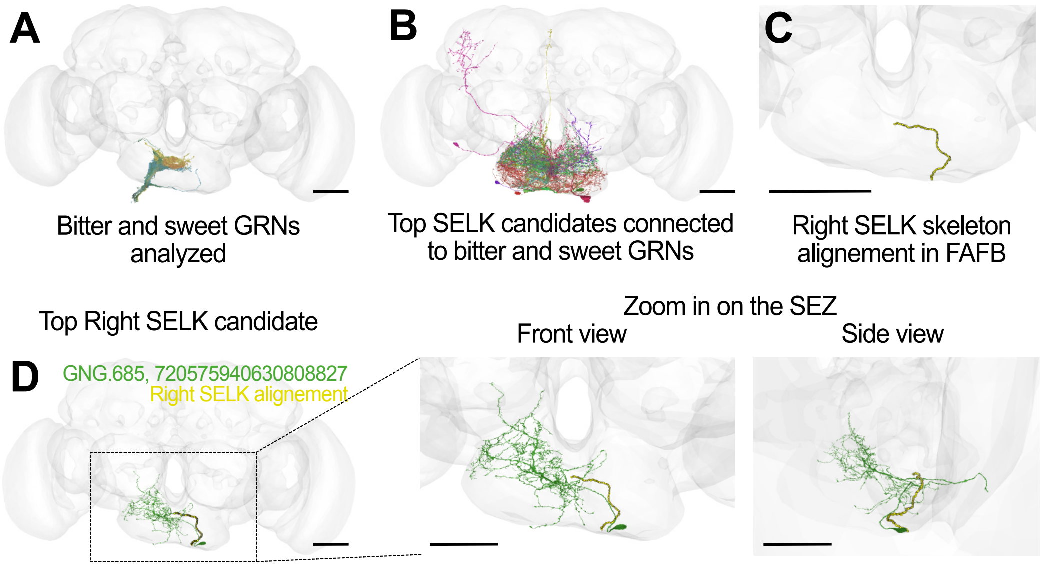


**Supplementary Figure 6. Sequential methodology to identify a strong SELK candidate neuron in the FAFB connectome.** (**A**) Reconstruction representation in the Full Adult Fly Brain (FAFB) brainmesh template in Flywire from all groups of GRNs analyzed, both bitter (group 1: orange and group 2: yellow) and sweet (group 4: green and group 5: blue) GRNs. (**B**) Top postsynaptic neurons to bitter and sweet GRNs analyzed considered as top SELK candidate neurons based on visual comparisons. (**C**) Alignment of the right SELK neuron segmented skeleton in the FAFB by using the Flywire Gateway app. (**D**) Top right SELK candidate neuron based on localization and morphology similarities with the right SELK neuron alignment from (**C**). ID from Flywire is indicated in green: 720575940630808827 (GNG.685/DNg68(R)). Zoom in on the SEZ from (**D**), showing detailed similarities of the right SELK candidate neuron arborization with the right SELK skeleton alignment both in front and side view. Scale bar: 50 µm.

**Supplementary Figure 7**

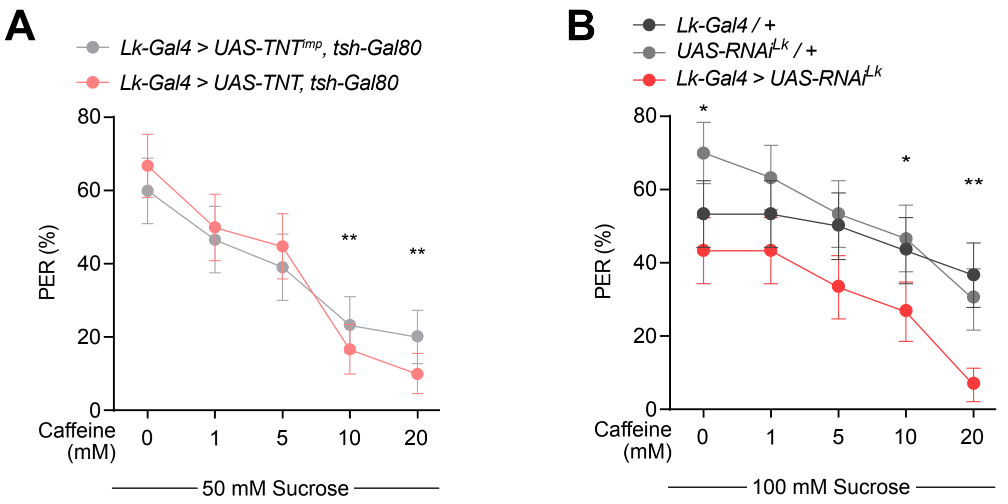


**Supplementary Figure 7. Leucokinin neurons are essential to discriminate sweet and bitter stimuli. (A)** PER of *Lk-Gal4 > tsh-Gal80, UAS-TNT^imp^* (n=30) and *Lk-Gal4 > tsh-Gal80, UAS-TNT* (n=30) for a for a mixture of 50 mM Sucrose with increasing concentrations of caffeine. (**B**) PER of *Lk-Gal4/+ , UAS-RNAi^Lk^/+ and Lk-Gal4 > UAS-RNAi^Lk^* (n=30), for a mixture of 100 mM Sucrose with increasing concentrations of caffeine. *p<0.05, **p<0.01. Logistic regression model with binomial distribution. Error bars represent the standard error of the proportion.
